# Spontaneous generation of innate number sense in untrained deep neural networks

**DOI:** 10.1101/857482

**Authors:** Gwangsu Kim, Jaeson Jang, Seungdae Baek, Min Song, Se-Bum Paik

**Affiliations:** Department of Physics, Korea Advanced Institute of Science and Technology, Daejeon 34141, Republic of Korea; Department of Bio and Brain Engineering, Korea Advanced Institute of Science and Technology, Daejeon 34141, Republic of Korea; Program of Brain and Cognitive Engineering, Korea Advanced Institute of Science and Technology, Daejeon 34141, Republic of Korea

## Abstract

Number-selective neurons are observed in numerically naïve animals, but it was not understood how this innate function emerges in the brain. Here, we show that neurons tuned to numbers can arise in random feedforward networks, even in the complete absence of learning. Using a biologically inspired deep neural network, we found that number tuning arises in three cases of networks: one trained to non-numerical natural images, one randomized after trained, and one never trained. Number-tuned neurons showed characteristics that were observed in the brain following the Weber-Fechner law. These neurons suddenly vanished when the feedforward weight variation decreased to a certain level. These results suggest that number tuning can develop from the statistical variation of bottom-up projections in the visual pathway, initializing innate number sense.

## Introduction

Number sense, an ability to estimate numbers without counting (Burr and Ross, 2008; Burr et al., 2018; Halberda et al., 2008; Piazza et al., 2010), is an essential function of the brain that may provide a foundation for complicated information processing (Nieder, 2016). It has been reported that this capacity is observed in humans and various animals in the absence of learning. Newborn human infants can respond to abstract numerical quantities across different modalities and formats (Feigenson et al., 2004; Izard et al., 2009; Xu et al., 2005), and newborn chicks can discriminate quantities of visual stimuli without training (Rugani et al., 2008, 2015). These results suggest that number sense arises spontaneously in the very early stages of the development.

In single-neuron recordings in numerically naïve monkeys (Viswanathan and Nieder, 2013) and crows (Wagener et al., 2018), it was observed that individual neurons in the prefrontal cortex and other brain areas can respond selectively to the number of visual items (numerosity). These results suggest that number-selective neurons (number neurons) arise spontaneously in the brain before visual experience and that they provide a foundation for innate number sense in young animals and humans. However, details of how number neurons originate are not yet understood.

Model studies with biologically inspired artificial neural networks have provided insight into the underlying mechanisms of brain functions, particularly with regard to the development of various functional circuits for visual information processing (Cadieu et al., 2014; Krizhevsky et al., 2012; Simonyan and Zisserman, 2015; Ullman et al., 2002; Yamins et al., 2014). Studies of number sense using deep neural networks (DNNs) have suggested that number neurons can emerge due to supervised and unsupervised learning with networks (Dehaene and Changeux, 1993; Stoianov and Zorzi, 2012; Verguts and Fias, 2004). A recent study showed that number-detector neurons are found in DNNs that were trained only for natural images, suggesting that number sense is an emergent function from training for visual object recognition (Nasr et al., 2019).

Here, we show that number tuning of neurons can spontaneously arise in completely untrained DNNs, enabling the network to perform a number discrimination task. Using a model network designed from the structure of a biological visual pathway, we found that number-selective neurons are observed in randomly initialized DNNs as well as in those trained with natural images. The responses of these neurons enabled the network to perform a number comparison task in this case. We also found that these number neurons show single- and multi-neuron characteristics that were observed in the brain following the Weber-Fechner law. From further investigations, we found that number neurons can originate solely from the weight variation of feedforward projections. Our findings suggest that innate number sense originates from number-tuned neurons that emerge spontaneously from the physical wiring of circuits during the early development stage.

## Results

### Emergence of number selectivity in networks trained for object classification

Number-selective neurons have been observed in various species, the response of which is maximized for a stimulus encoding a specific numerosity (**Fig. 1A**) (human: (Kutter et al., 2018), monkey: (Nieder and Merten, 2007), crow: (Wagener et al., 2018)). One important feature observed in these neuronal responses is the Weber-Fechner law, where the width (*σ*, sigma of the Gaussian fit) of the tuning curves increases proportionally with an increase in the numerosity on a linear scale (**Fig. 1B**, left top and orange dots in **1B**, right; slope = 0.38), or equivalently remains constant across the numerosity on a logarithmic scale (**Fig. 1B**, left bottom and green dots in **1B**, right; slope = −0.0091) (Nieder and Merten, 2007).

**Fig 1.**
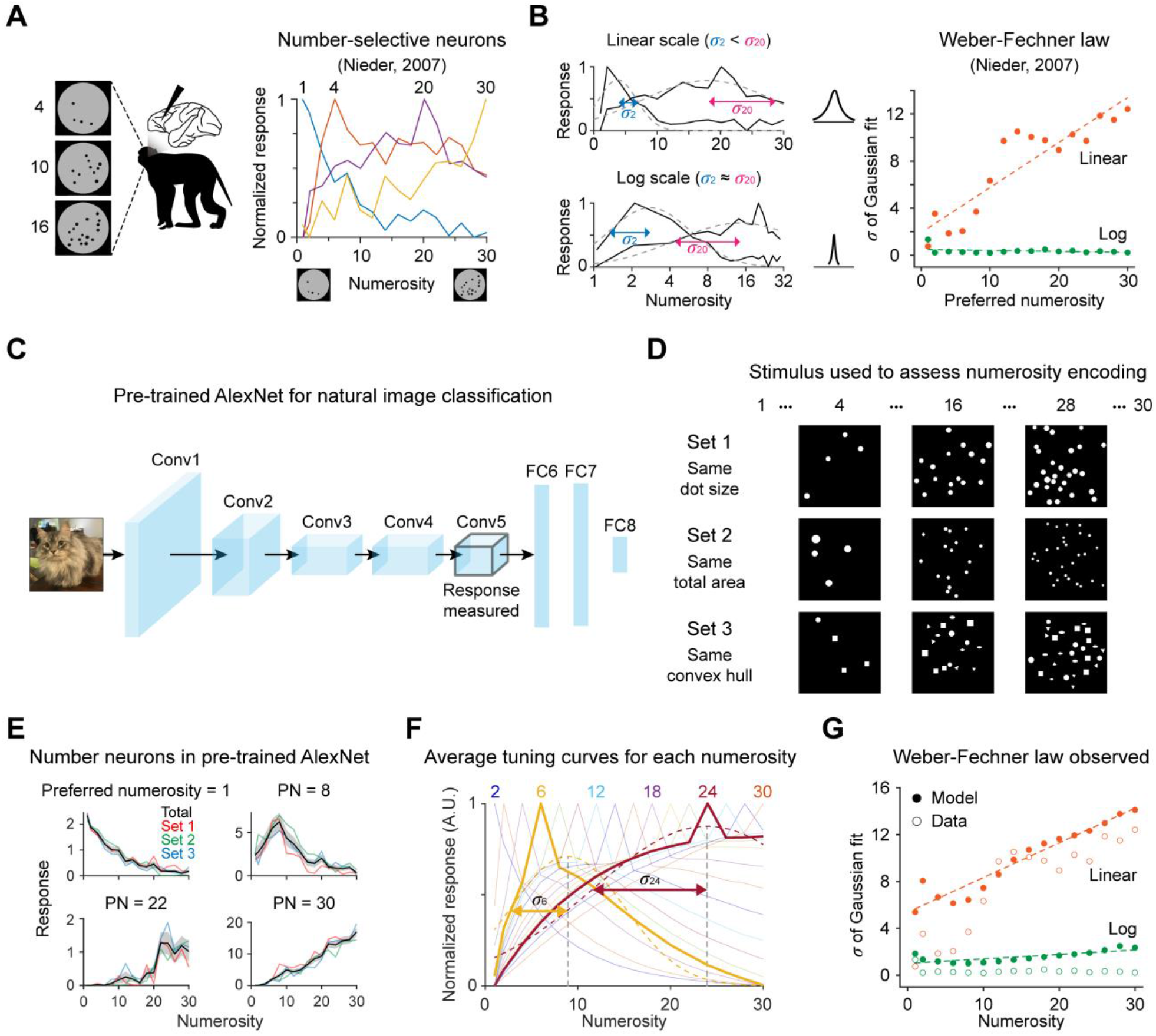
Emergence of number selectivity in networks trained for object classification. (A) Number-selective neurons observed in monkeys (Nieder and Merten, 2007) (B) The observed neuronal number tuning is described by the Weber-Fechner law such that the tuning width increases proportionally with an increase in the numerosity on a linear scale (left top and orange dots on the right) or remains constant across the numerosity on a logarithmic scale (left bottom and green dots on the right). Left, Dashed lines indicate a fitted Gaussian distribution. Right, Dashed lines indicate linear fitting. (C) Architecture of the pre-trained AlexNet. (D) Examples of the stimuli used to measure number tuning (Nasr et al., 2019). Set 1 contains dots of the same size. Set 2 contains dots with a constant total area. Set 3 contains items of different geometric shapes with an equal overall convex hull. (E) Examples of tuning curves of individual number-selective neurons as reported (Nasr et al., 2019). (F) Average tuning curves of different numerosities on a linear scale. Note that the tuning width (sigma of the Gaussian fitting) increases as the preferred numerosity increases. (G) The tuning width increases proportionally on a linear scale and remains constant on a logarithmic scale, as predicted by the Weber-Fechner law.

We initially reproduced previous results indicating that number-selective neurons emerge in a deep neural network trained for object classification (Nasr et al., 2019) (**Figs. 1C-G**). To explore the developmental origin of these number neurons, we simulated the response of AlexNet, a conventional deep neural network that models the ventral visual stream of the brain (**Fig. 1C**) (Krizhevsky et al., 2012). The network consists of five convolutional layers for feature extraction and three fully connected layers for object classification. In the current study, to investigate the selective responses of neurons rather than the performance of the system, the classification layers were discarded and the responses of units in the last convolutional layer (conv5) were examined.

From a simulation of this model network, we confirmed the observed results of Nasr et al. that number-selective neurons emerge from the training of the network for object classification. Using a pre-trained AlexNet (Krizhevsky et al., 2012) (**Fig. 1C**) — trained for the classification of natural images from the ILSVRC2010 ImageNet database — we tested the number-selective responses of neurons for images of dot patterns depicting numbers spanning 1 to 30 (Nieder and Merten, 2007) (**Fig. 1D**). With a test design introduced in earlier work (Nasr et al., 2019), we used three different sets of stimuli to ensure the invariance of the observed number tuning for certain geometric factors (Zhaoping, 2002), in this case the stimulus size, density, and area (set 1, circular dots of the same size; set 2, dots equal the total dot area; set 3, items of different geometric shapes with an equal overall convex hull). As a result, we confirmed that neurons tuned to various instances of numerosity were observed as reported (**Fig. 1E**). We also confirmed that these neurons satisfy the Weber-Fechner law, as observed in the neurons in the brain (**Figs. 1F** and **1G**; linear, slope = 0.30; log, slope = 0.038).

### Spontaneous emergence of number-selective neurons in untrained networks

Subsequently, to examine whether the training process for object classification is essential for the emergence of number-selective units, we devised an untrained AlexNet by randomly permuting the weights of filters in each convolutional layer (**Fig. 2A**; permuted AlexNet) such that the network loses its ability to classify objects (**Fig. 2B**; *p* = 3.00×10^−37^, Wilcoxon rank-sum test). Surprisingly, we found that number-selective neurons were still observed in the permuted AlexNet despite the fact that the network was never trained with visual stimuli after the randomization step (**Fig. 2C** and **Supplementary Fig. 1**; 9.58% of whole units). The sharpness of tuning in individual units appears to be slightly broader than that in the trained network (**Supplementary Fig. 2**; goodness of the Gaussian fit, *r*^2^ = 0.77±0.19 in the pre-trained case; 0.72±0.21 in the permuted case).

**Fig 2.**
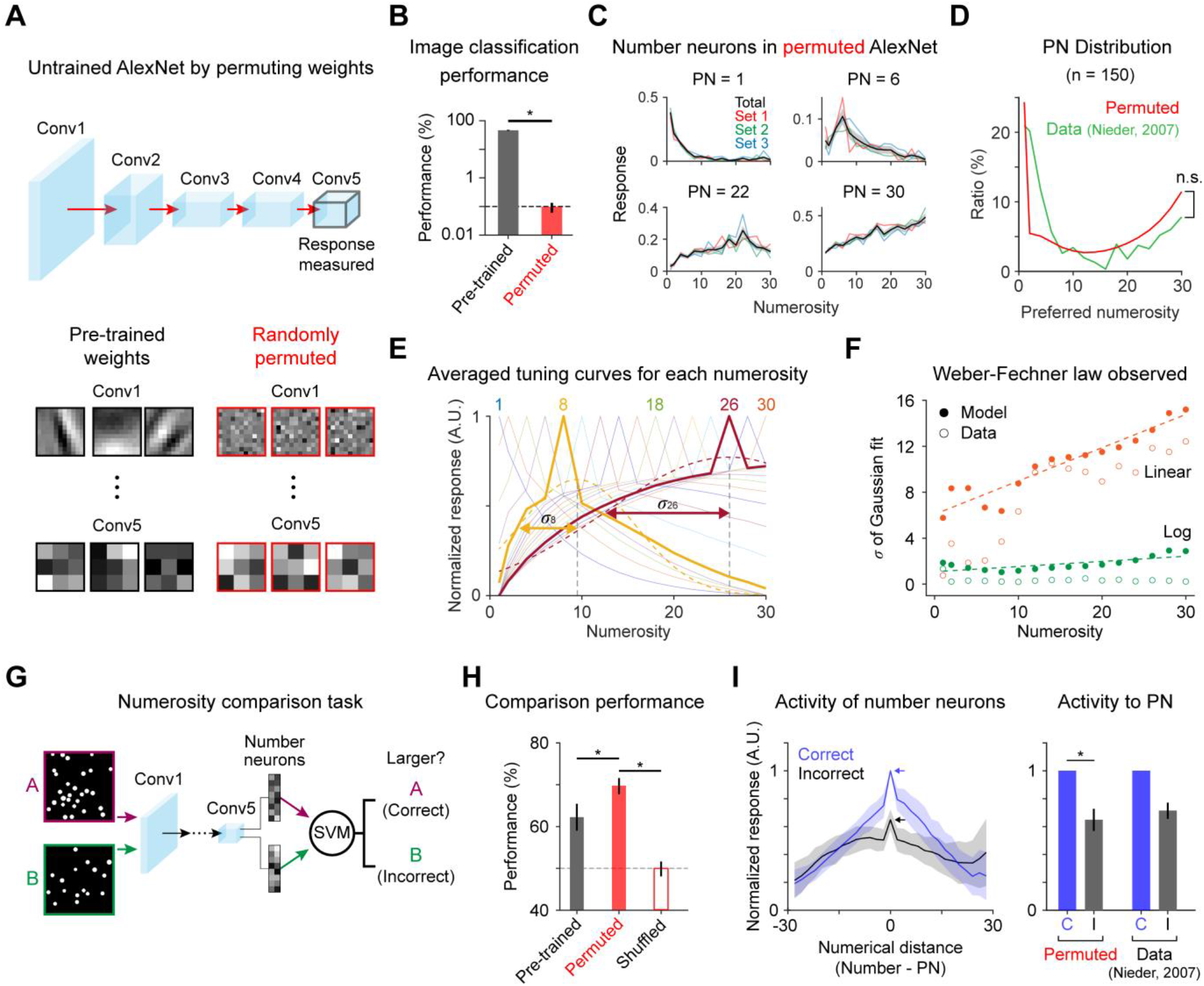
Spontaneous emergence of number selectivity in untrained networks. (A) An untrained AlexNet was devised by randomly permuting the weights in each convolutional layer of a trained network. (B) The permuted network loses its object classification ability. The dashed line indicates the chance level (0.1%). *p* = 3.00× 10^−37^; Wilcoxon rank-sum test. (C) Examples of tuning curves for individual number-selective network units. (D) Distribution of preferred numerosity in the network and the observation in monkeys (Nieder and Merten, 2007). The root mean square error between the red and green curves (permuted vs. data) is significantly lower than that in the control with the shuffled distribution (*p* = 0.01). (E) Average tuning curves of different numerosities on a linear scale. (F) The tuning width increases proportionally on a linear scale and remains constant on a logarithmic scale, as predicted by the Weber-Fechner law (Nieder and Merten, 2007). (G) Number comparison task using the SVM. (H) Task performance of the permuted network, pre-trained network and the control of shuffled responses: permuted vs. control (red solid vs. red open bar), *p* = 2.56×10^−34^; pre-trained vs. control (dark gray solid vs. red open bar), *p* = 2.56× 10^−34^; permuted vs. pre-trained (red solid vs. dark gray solid bar), *p* = 2.56× 10^−34^, Wilcoxon rank-sum test. (I) Left, Average activity of number-selective units as a function of the numerical distance. Right, Response to the preferred numerosity; note that the response during correct trials is significantly higher than that during incorrect trials, as observed in actual neurons recorded from a monkey prefrontal cortex during a numerosity matching task (Nieder and Merten, 2007). *p* = 5.64×10^−39^, Wilcoxon rank-sum test.

Previously observed profiles of tuned neurons in monkeys (Nieder and Merten, 2007), including the Weber-Fechner law, were also reproduced by number neurons in the permuted AlexNet (**Figs. 2D-F**). First, the distribution of the preferred numerosity covered the entire range (1~30) of the presented numerosity (**Fig. 2D**, red curve). Moreover, number neurons preferring 1 or 30 were most frequently observed such that the ratio of neurons increases as the preferred numerosity decreases to 1 or increases to 30, with a profile similar to the experimental observation in monkeys (**Fig. 2D**, green curve) (Nieder and Merten, 2007). In addition, as the numerosity increases, the sigma of the Gaussian fit of the averaged tuning curve increased on a linear scale (**Fig. 2E**), following the Weber-Fechner law, as observed in biological brains. (**Fig. 2F**; linear, slope = 0.29; log, slope = 0.044).

Next, we examined if these number-selective neurons can perform a number comparison task (**Fig. 2G**; see Methods for details). In the task, two images of dot patterns were presented and the network determined which stimulus has greater numerosity from the output of a support vector machine (SVM) trained with the responses of number-selective neurons. The measured correct performance rate of the network was found to be 69.7±1.9% (**Fig. 2H**, red solid bar), even slightly better than that in the pre-trained network (62.2±3.2%; **Fig. 2H**, dark gray solid bar) and significantly higher than that when the responses of the neurons were shuffled across two presented images (**Fig. 2H**, red open bar; permuted vs. control (red solid vs. red open bar), *p* = 2.56×10^−34^; pre-trained vs. control (dark gray solid vs. red open bar), *p* = 2.56×10^−34^; permuted vs. pre-trained (red solid vs. dark gray solid bar), *p* = 2.56×10^−34^, Wilcoxon rank-sum test). This result suggests that the selective responses of observed number neurons can provide the network with the capability to compare numbers. To investigate the contributions of the number-selective neurons for correct choices in greater depth, we compared the average tuning curves obtained in correct and incorrect trials (**Fig. 2I**) (Nasr et al., 2019). As expected, the average response to the preferred numerosity in incorrect trials significantly decreased to 64.8% of that in correct trials (**Fig. 2I**; blue arrow vs. black arrow; *p* = 5.64×10^−39^, Wilcoxon rank-sum test), as observed in the numerosity matching task with monkeys (**Fig. 2I**, right) (Nieder and Merten, 2007).

### Number-selective neurons from the weighted sum of increasing and decreasing feedforward activities

Subsequently, we examined how number-selective neurons emerge in untrained random feedforward networks. Important clues were found in the neurons observed in all layers (Conv1-Conv5), the responses of which monotonically decrease or increase as the stimulus numerosity increases (**Fig. 3A**, decreasing and increasing units), as suggested by previous modelling studies (Dehaene and Changeux, 1993; Verguts and Fias, 2004). Notably, we found that decreasing units (N = 785 ± 208 in Conv4; 1.21 ± 0.32% of all units) showed nonzero responses to blank inputs (**Fig. 3A**, top, blank response), with their responses decreasing from the blank response as the numerosity in the stimulus increased (**Fig. 3A**, top). In contrast, we observed that increasing units (N = 1,474 ± 217 in Conv4; 2.27 ± 0.33% of all units) showed blank responses close to zero, with their responses increasing from that level as the numerosity increased (**Fig. 3A**, bottom).

**Fig 3.**
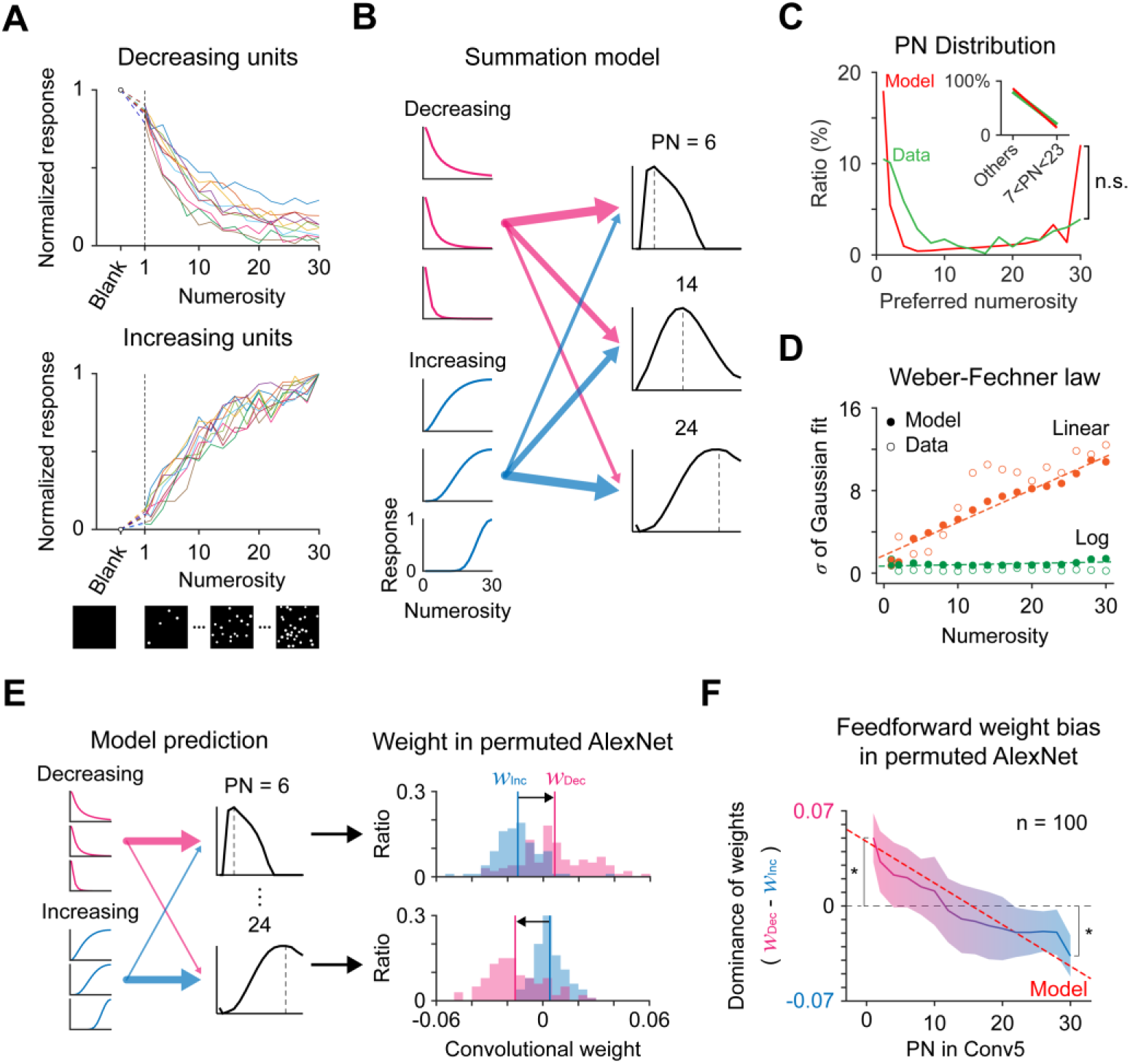
Emergence of number tuning to diverse values of numerosity from the weighted sum of increasing and decreasing unit activities. (A) Instances of monotonically decreasing or increasing neuronal activity as the numerosity of the stimulus increases were observed in all layers (Conv1-Conv5). (B) Number tuning as the summation of decreasing and increasing units. Neurons tuned to various numbers can be generated from the weighted sum of increasing and decreasing tuning curves. (C) Distributions of the preferred numerosity from the summation model and observations in monkeys (Nieder and Merten, 2007). The root mean square error between the red and green curves (simulation vs. data) is significantly lower than that in the control with the shuffled distribution of the simulated case (*p* = 0.01). (D) The tuning width increases proportionally on a linear scale and remains constant on a logarithmic scale, as predicted by the Weber-Fechner law (Nieder and Merten, 2007). (E) Left, a summation model of tuning curves to various preferred numbers developed from a linear combination of increasing and decreasing unit activities. Neurons tuned to smaller numbers receive strong inputs from the decreasing units and receive weak inputs from the increasing units, while neurons tuned to larger numbers receive stronger inputs from increasing units. Right, bias in the convolutional weights predicted by the model. (F) The weight bias of all number neurons observed in conv5 of the permuted AlexNet. As predicted by the summation model (red dashed line), the neurons tuned to smaller numbers receive stronger inputs from decreasing units, and vice versa. *p* = 3.90×10^−18^ at PN = 1; *p* = 4.27×10^−18^ at PN = 30, Wilcoxon signed-rank test.

We hypothesized that these increasing and decreasing units can be the building blocks of number-selective neurons tuned to various numerosity values. We developed a summation model which holds that tuning curves tuned to various preferred numbers develop from a linear combination of increasing and decreasing unit activities (modeled as lognormal distributions peaking at 1 and 30, respectively) (**Fig. 3B**). A simulation of this simple model showed that number-selective neurons of all PNs can arise from the weighted sum such that neurons tuned to lower numbers receive strong inputs from decreasing units and receive weak inputs from increasing units, while neurons tuned to higher numbers receive stronger inputs from increasing units. From 200,000 repeated trials in which the weights of feedforward linear combinations were randomized, we found that number-selective neurons preferring 1 or 30 were most frequently observed such that the ratio of tuned neurons increases as the preferred numerosity decreases to 1 or increases to 30 (**Fig. 3C**). This result implies that tuning to various numerosity values originates from the weighted sum of the decreasing units (PN = 1) and increasing units (PN = 30). In this model, we also confirmed that the sigma of the Gaussian fit of the averaged tuning curve increased on a linear scale (**Fig. 3D**) as the numerosity increases, following the Weber-Fechner law.

Next, we examined whether the number-selective neurons observed in conv5 of the permuted AlexNet show expected bias in feedforward input weights between decreasing and increasing units (**Fig. 3E**, left). We measured all connections from decreasing and increasing units in the previous layer and compared the average weights from decreasing units with those from the increasing units (**Fig. 3E**, right). As predicted, neurons tuned to smaller numbers received stronger inputs from decreasing units while neurons tuned to larger numbers received stronger inputs from increasing units (**Fig. 3F**), implying that the observed neuronal tuning for various levels of numerosity originated from the summation of the monotonically decreasing and increasing activity units in the earlier layers.

### Number selectivity by diversity of the convolutional weight

To examine the origin of number tuning in untrained neural networks, we implemented a randomly initialized network (randomized AlexNet) (**Fig. 4A**). In this model, the values of each weight kernel were randomly sampled from a Gaussian distribution that fits the weight distribution of the pre-trained network, with weight variation of the feedforward kernels be controllable by modulating the width of the Gaussian. Using this model network, we investigated if the emergence of number selectivity is affected by the statistical variation of random feedforward projection.

**Fig 4.**
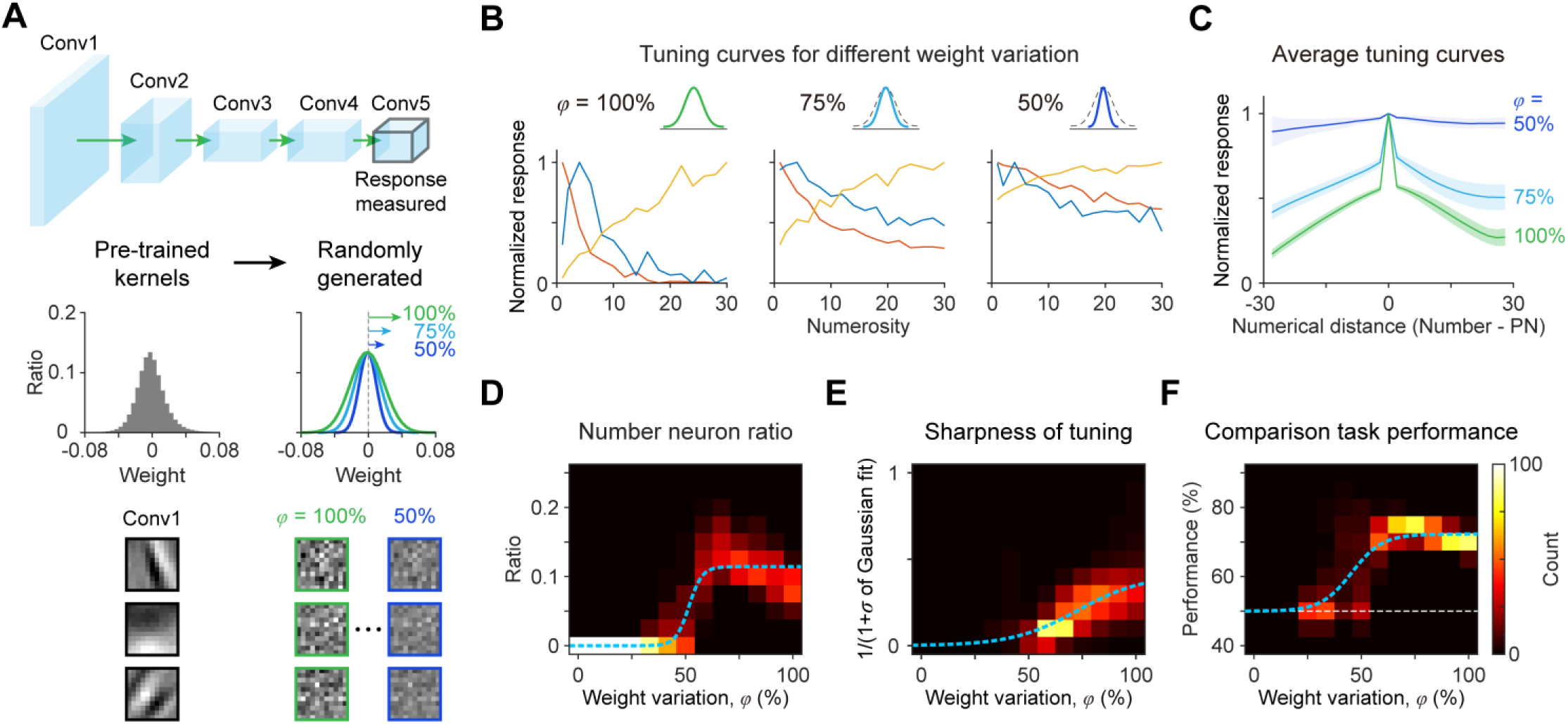
Number tuning induced by variations in convolutional weights. (A) An untrained AlexNet was generated from the random sampling of values in each weight kernel from a Gaussian distribution that fits the weight distribution of the pre-trained condition. The weight variation can be controlled by modulating the standard deviation of the Gaussian. Changes of tuning curves for individual number-selective neurons across different levels of weight variation. (C) Average of all tuning curves. Note that tuning becomes broader as the weight variation is reduced. (D) When the weight variation is reduced to less than 70% of that in the pre-trained network, the ratio of number neurons is suddenly decreased. (E) The sharpness of tuning decreases as the weight variation is reduced. The ratio of units for each weight variation factor (*φ*) was normalized to the number of all number-selective units at *φ* = 100%. (F) As the weight variation is reduced, the performance on the numerosity comparison task decreases to the chance level (white dashed line). (D-F) Blue dashed lines indicate fitting for the sigmoid function.

First, we observed that the randomized AlexNet with weight variations equivalent to the pre-trained conditions (weight variation factor, *φ* = 100%) also generated neurons tuned for numerosity, as observed previously (**Fig. 4B**, left). We then gradually reduced the weight variation of each kernel and examined changes in number tuning of the units. When the weight variation decreased to half (*φ* = 50%) of the original value, number-selective neurons were observed more rarely (3.90% of all units = 42.1% of selective neurons at *φ* = 100%) and the average tuning curve of the neurons became nearly flat (**Fig. 4B**, right and **4C**). Importantly, we found that the number of selective neurons suddenly decreased when the weight variation was reduced to less than 70% of that in the pre-trained network (**Fig. 4D**; *r*^2^ of the fit for the sigmoid function (*ƒ*(*x*) = *a*/(1 + *b*exp(−*cx*)) = 0.81, *r* = 0.77, *p* < 10^−40^). Moreover, the sharpness of tuning (1/(1+*σ* of the Gaussian fit)) and the SVM performance during the number comparison task also decreased monotonically as the weight variation was reduced (**Figs. 4E** and **4F**; *r*^2^ of the fit for the sigmoid function = 0.36 and 0.62, *r* = 0.58 and 0.66, *p* < 10^−40^ and *p* < 10^−40^, respectively), suggesting that a certain level of variation in the convolutional weights is required for the spontaneous emergence of number tuning. These results imply that innate number neurons develop solely from the statistical variation in the random initial wirings of bottom-up projections in the visual pathway.

## Discussion

Using biologically inspired deep neural network models, we showed that number neurons can spontaneously emerge in a randomly initialized network without any learning. We found that a statistical variation of the weights in feedforward projections is a key factor to generate neurons tuned to numbers. These results suggest that neuronal tunings, which initialize innate functions of the brain, arise exclusively from the statistical complexity embedded in the neural circuitry.

These findings provide an explanation of how number neurons or number sense similarly arise in a wide range of different species, from mammalian (Viswanathan and Nieder, 2013) to avian (Wagener et al., 2018), separated in the phylogenetic tree by nearly 300 million years ago (Cardew and Bock, 2000; Ditz and Nieder, 2015). In particular, observations of number neurons in these naïve animals of various species suggest that number tuning emerges regardless of the species-specific design of the neural circuitry, such as different numbers of layers in the neocortex. Instead, our findings suggest that number neurons can originate from much simpler and more common components in the feedforward afferents. This may provide the simplest theoretical scenario for the spontaneous emergence of number neurons at the early developmental stage in various species with different organizations of the cortical circuits.

Next, the emergence of number sense in a random feedforward network may provide an expanded framework with which to understanding other innate visual functions in the brain (Ullman et al., 2012). In recent studies, it was suggested that various cognitive functions can also emerge spontaneously in randomized neural networks: it was argued that selective tunings such as number-selective responses may be generated simply from the multiplication of random matrices (Hannagan et al., 2018). It was also shown that the structure of a randomly initialized convolutional neural network can provide a priori information about the low-level statistics in natural images, enabling the reconstruction of the corrupted images without any training for feature extraction (Ulyanov et al., 2018). Gestalt phenomena in human visual perception was modeled in an untrained neural network, with the results implying the ability of a randomly initialized network to engage in visual feature extraction (Kim et al., 2019). A generalized model of a randomly connected neural network for extracting higher order features of inputs has been actively studied in relation to reservoir computing (Lukoševičius and Jaeger, 2009; Tanaka et al., 2019). Overall, these results imply that the random initial states of a neural network during early development are able to provide a template for various innate functions of the brain with zero-shot training for visual tasks (Palatucci et al., 2009; Zhang et al., 2018).

Lastly, the question arises of how the results of the DNN models in the current study can be compared with the development of early cortical circuits in the brain. In the early development stage before visual experience, the visual pathway from the retina to the cortex is initialized by retinotopic projections of feedforward afferents (Jang and Paik, 2017; Paik and Ringach, 2012, 2011; Ringach, 2004, 2007; Sailamul et al., 2017). During this stage, local wirings for feedforward convergence projection are noisy and the receptive fields of individual neurons are not yet refined (Sarnaik et al., 2014; Tavazoie and Reid, 2000). This is comparable to the condition of a convolutional filter in a randomly initialized neural network before training for a task. In this stage of DNNs, a convolutional process is simply a local sampling of feedforward inputs with random weights. It was previously believed that convolutional filters must be refined by training for the network to perform a function, but here we suggest that the network can perform certain innate functions with these untrained filters. In the case of a biological neural network, spontaneous tunings generated in this early condition will initialize various functions, possibly leading to highly effective refinement of the receptive fields when learning begins with sensory inputs.

In summary, we conclude that the innate number tuning of neurons can spontaneously arise in a completely untrained neural network, solely from the statistical variance of feedforward projections. This finding suggests that various innate functions in the brain may originate from the organization of the random initial wirings of neural circuits, thus providing new insight into the origin of innate cognitive functions.

## Methods

### Neural network model

AlexNet (Krizhevsky et al., 2012) was used as a representative model of a convolutional neural network. It consists of five convolutional layers with rectified linear unit (ReLU) activation, followed by three fully connected layers. The detailed designs and hyperparameters of the model were determined based on earlier work (Krizhevsky et al., 2012).

Based on the architecture described above, three types of variations of the network (pre-trained, permuted, and randomized) were investigated. Pre-trained AlexNet: The network was trained to classify objects in the ImageNet dataset (Russakovsky et al., 2015) such that the model parameters including the weights of convolutional filters and biases were refined to extract the high-level features of natural images. The pre-trained AlexNet model was obtained from the MATLAB Deep Learning Toolbox. Permuted AlexNet: For each layer, the values of the weights and biases were randomly permuted such that the overall distribution is preserved while any spatial patterns in any kernel are removed. Randomized AlexNet: For each layer, the mean and the standard deviation of the distribution of weights (and biases) were calculated. Then, each weight (and bias) was re-determined by a Gaussian distribution having the same mean and the same (or reduced) standard deviation. The ability of networks to classify objects was tested with 1,000 images of different classes, which were randomly chosen from the ILSVRC2010 ImageNet database. All simulations underwent 100 trials.

### Stimulus dataset

The stimulus sets used in this work were designed based on earlier work (Nasr et al., 2019). Briefly, images (size: 227 × 227 pixels) that contain N = 1, 2, 4, 6, …, 28, 30 circles were provided as inputs to the network. To ensure the invariance of the observed number tuning for geometric factors such as the stimulus size, density, and area, three stimulus sets were designed. In set 1, dots were located at random locations (without overlapping) but with a nearly consistent radius (generated by the normal distribution; mean = 7, STD = 0.7). In set 2, the total area of the dots remains constant (1,200 pixel^2^) across different numerosities, and the average distance between neighboring dots is constrained in a narrow range (90 – 100 pixels). In set 3, a convex hull of the dots was fixed as a regular pentagon, the circumference of which is 647 pixels. The shape of each dot was determined to be that of a circle, a rectangle, an ellipse, and a triangle with an equal probability for each. Fifty images were generated for each combination of the numerosity and the stimulus set, meaning that 50×16×3 = 2400 images were used in total to evaluate the responses of the network units.

### Analysis of the responses of the network units

The responses of network units in the final convolutional layer (after ReLU activation of the fifth convolutional layer) were analyzed. Similar to the method used to find number-selective neurons in monkeys (Nieder and Merten, 2007) and to detect number-selective network units (Nasr et al., 2019), a two-way ANOVA with two factors (numerosity and stimulus set) was used. To detect number-selective units generating a significant change of the response across numerosities but with an invariant response across stimulus sets, a network unit was considered to be number-selective if it exhibited a significant effect for numerosity (*p* < 0.01) but no significant effect for the stimulus set or interaction between two factors (*p* > 0.01). The preferred numerosity of a unit was defined as the numerosity that induced the largest response on average among the responses for all presentations. The tuning width of each unit was defined as the sigma of the Gaussian fit of the average tuning curve on a logarithmic number scale.

To determine the average tuning curves of all number-selective units, the tuning curve of each unit was normalized by mapping the maximized response to 1 and was then averaged across units using the preferred numerosity value as a reference point. To compare the average tuning curves across different numerosities, the tuning curve of each unit was averaged across units preferring the same numerosity and was then normalized by mapping the minimized and maximized responses to 0 and 1, respectively. In the analysis with the reduced weight variations, the units with an average response weaker than 0.05 times that in the randomized AlexNet (*φ* = 100%) were not considered in the analysis so as to ignore cases where a weak fluctuation of the response was measured as a number-selective case.

### Numerosity comparison task for the network

A numerosity comparison task was designed to examine whether number-selective neurons can sufficiently perform a numerical task that requires an estimation of numerosity from images. For each trial, a sample and a test stimulus (randomly selected from 1, 2, 4, …, 28, 30) were presented to the network and the resulting responses of the number-selective neurons were recorded. Then, a support vector machine (SVM) was trained with the responses of 16 randomly chosen number-selective neurons to predict whether the numerosity of the sample stimulus is greater than that of the test stimulus. In this case, 100 sample stimuli were generated for each form of numerosity (1600 stimuli in total) and the test stimuli for each sample were generated while avoiding the numerosity of the corresponding sample stimulus. To calculate the average tuning curves for the correct and incorrect trials, the tuning curve of each neuron was normalized so that response for the preferred numerosity in the correct trial was mapped to 1.

---

**Supplementary Figure 1.**
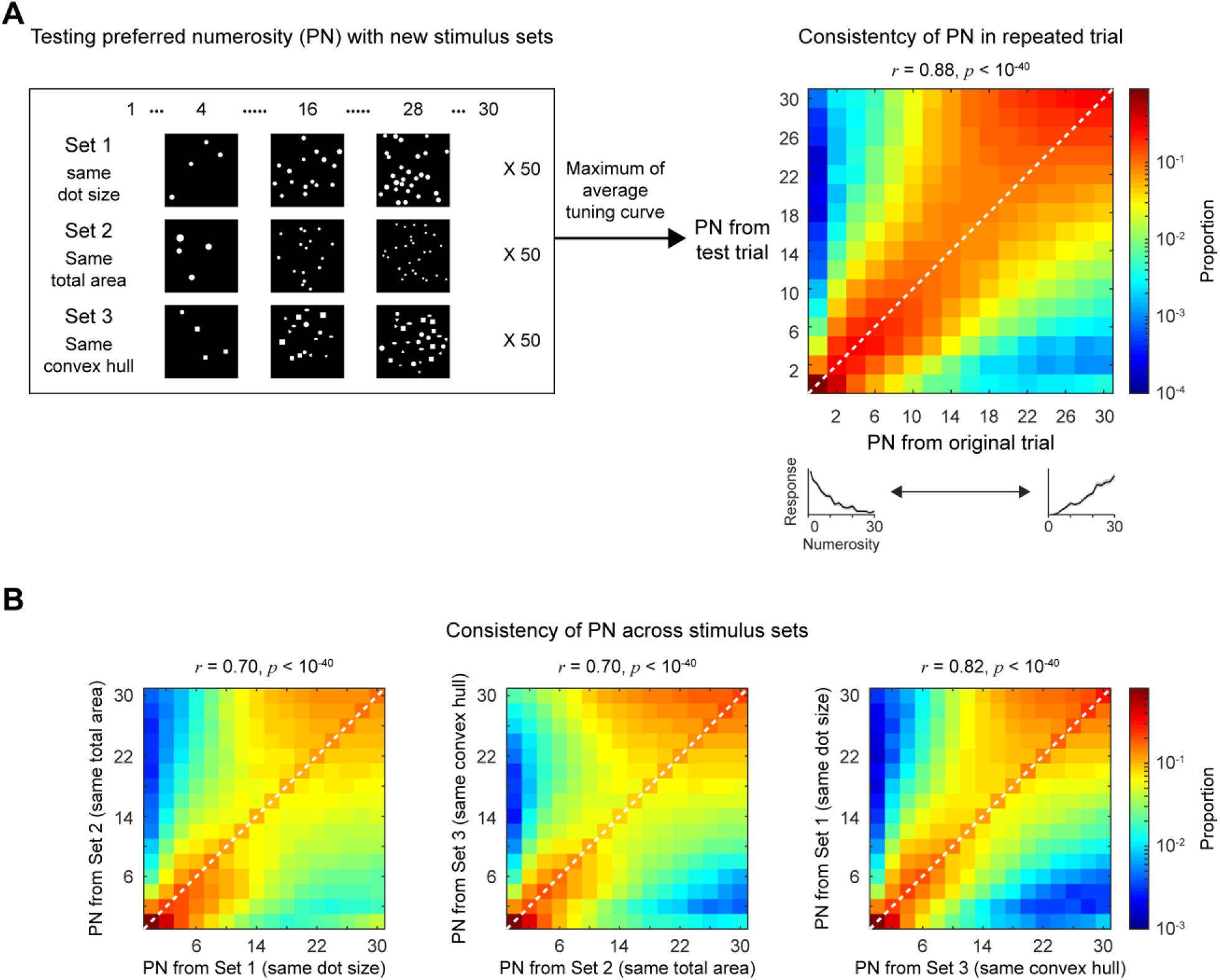
Consistency of the preferred numerosity of number neurons. The preferred numerosity (PN) outcomes measured with different stimuli are significantly correlated with each other, implying consistency of the preferred numerosity. (A) PN measured with the original stimulus sets vs. PN measured with newly generated stimulus sets. (B) PNs measured with each type of stimulus condition were compared.

**Supplementary Figure 2.**
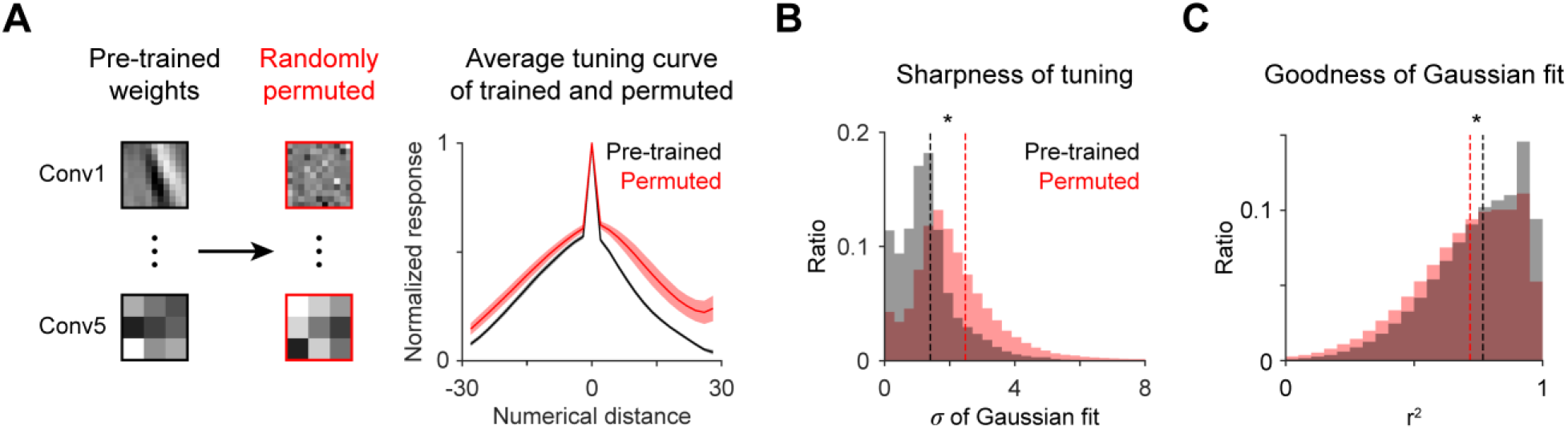
Sharpness of tuning appears to be slightly broader in the permuted AlexNet. (A) Average tuning curves obtained from the pre-trained and the permuted AlexNet. (B-C) The sharpness of tuning (B) and the goodness of the Gaussian fit (C) in individual units of the permuted network respectively appear to be slightly broader and better fitted than that in the pre-trained network. *p* < 10^−40^; Wilcoxon rank-sum test

